# Stimulus-induced narrowband gamma oscillations are test-retest reliable in healthy elderly in human EEG

**DOI:** 10.1101/2021.07.06.451226

**Authors:** Wupadrasta Santosh Kumar, Keerthana Manikandan, Dinavahi V.P.S. Murty, Ranjini Garani Ramesh, Simran Purokayastha, Mahendra Javali, Naren Prahalada Rao, Supratim Ray

## Abstract

Visual stimulus-induced narrowband gamma oscillations in electroencephalogram (EEG) recordings have been recently shown to be compromised in subjects with Mild Cognitive Impairment or Alzheimer’s Disease (AD), suggesting that gamma could be an inexpensive and easily accessible biomarker for early diagnosis of AD. However, to use gamma as a biomarker, its characteristics should remain consistent across multiple recordings, even when separated over long intervals. Previous magnetoencephalography studies in young subjects have reported that gamma power remains consistent over recordings separated by a few weeks to months. Here, we assessed the consistency of slow (20-35 Hz) and fast gamma (36-66 Hz) oscillations induced by static full-field gratings in male (N=20) and female (N=20) elderly subjects (>49 years) in EEG recordings separated by more than a year, and tested the consistency in the magnitude of gamma power, its temporal evolution and spectral profile. Gamma oscillations had distinct spectral and temporal characteristics across subjects, which remained consistent across recordings (average intraclass correlation, ICC of ∼0.7). Alpha oscillations (8-12 Hz) and steady-state-visually-evoked-potentials (SSVEPs) were also found to be reliable. We further tested how EEG features can be used to identify two recordings as belonging to the same versus different subjects and found high classifier performance (area under ROC curve of ∼0.89), with the temporal evolution of slow gamma and spectral profile emerging as the most informative features. These results suggest that EEG gamma oscillations are reliable across recordings and can be used as a clinical biomarker as well as a potential tool for subject identification.

**Significance statement:** We demonstrate the reliability of stimulus-induced gamma oscillations in elderly humans for the first time in EEG. Since gamma has recently been shown to be compromised in patients with Mild Cognitive Impairment or early Alzheimer’s Disease (AD), together these results mark the first steps towards an EEG based clinical biomarker for early diagnosis of AD. We observed high reliability in the power spectrum, gamma power and its temporal characteristics, within the test-retest period of one year. Alpha and steady-state-visually-evoked potential power were also found to be reliable. These spectral and temporal features could also be used to identify EEG recordings as belonging to the same versus different subjects with high performance, suggesting a potentially key role in subject identification also.

## Introduction

Gamma rhythm corresponds to narrow-band oscillatory activity between 30-70 Hz in the brain, which has been shown to be involved in many higher cognitive functions like attention (Fries et al., 2001; Jensen et al., 2007) and memory (Herrmann et al., 2004). Gamma oscillations have also been shown to be affected in cognitive disorders such as Schizophrenia and Autism (Uhlhaas and Singer, 2010, 2012). In addition, gamma oscillations can be induced by presenting certain visual stimuli such as bars and gratings, and their amplitude and centre frequency dependent on the stimulus properties such as size, contrast, and spatial frequency (Henrie and Shapley, 2005; Gieselmann and Thiele, 2008; Ray and Maunsell, 2010; Jia et al., 2013; Murty et al., 2018). Recently, we have shown that these stimulus-induced narrowband gamma oscillations recorded using electroencephalogram (EEG) in elderly subjects (>49 years) weaken with age (Murty et al., 2020) and are weaker in subjects with Mild Cognitive Impairment (MCI) and early Alzheimer’s Disease (AD) compared to their age and gender matched controls (Murty et al., 2021). These results suggest that stimulus-induced gamma oscillations recorded using EEG could potentially be an inexpensive and easily accessible biomarker for diagnosis of mental disorders such as AD.

For the induced gamma to serve as a clinical biomarker, it is necessary to confirm its consistency across multiple recording sessions. Previous studies have shown that stimulus-induced visual gamma oscillations are test-retest reliable, but there are several limitations. First, such test-retest reliability of induced visual gamma oscillations is limited to magnetoencephalogram (MEG) recordings (Hoogenboom et al., 2006; Muthukumaraswamy et al., 2010; Tan et al., 2016). The only EEG study to our knowledge that reported induced gamma oscillations used a checkerboard stimulus that mainly induced broadband gamma (Keil et al., 2003); another study that used gratings to induce gamma oscillations reported reliability of evoked, not induced, gamma oscillations (Fründ et al., 2007). Second, although having a test-retest interval of a year allows for checking its feasibility as an annual check-up, most of these studies have tested reliability with test-retest interval spanning from a few weeks to months (but see McCusker et al., 2021 who had a test-retest interval of 3 years, although they used checkerboard patterns and their paradigm also included a behavioural task). Third, most studies have been conducted on young subjects (20-40 years). To test capability of gamma power as a clinical biomarker, it is important to consider the geriatric subjects who are prone to cognitive disorders. Further, steady-state-visually-evoked-potentials (SSVEPs) at gamma frequencies have recently been shown to have therapeutic effect on rodent models of AD (Iaccarano et al., 2016), and to be useful in Brain Computer Interfacing (BCI) applications (Vialatte et al., 2010). Therefore, it is important to compare the test-retest reliability of SSVEPs with induced gamma oscillations. Finally, most previous studies have focused on power and centre frequency of induced gamma oscillations. However, these oscillations also have a characteristic temporal evolution, and in addition, the peak frequencies of these and other oscillations vary across individuals, thus generating a characteristic spectral profile as well. Since these spectral and temporal features could provide additional information beyond simply the power and centre frequency, it is important to study test-retest reliability of these features. Such power and centre frequency based features have been studied in the context of subject identifiability (Näpflin et al., 2007) and biometric authentication (Thomas and Vinod, 2018) in resting state EEG data, but not for stimulus-induced gamma oscillations.

To address these limitations, we tested the reliability of visually induced narrowband gamma oscillations recorded using EEG in healthy elderly (aged > 49 years) with an interval of one year or more. This dataset is a part of double-blind case-control EEG study (N=257; 106 females), part of which has been used earlier to show that gamma oscillations weaken with healthy ageing (Murty et al., 2020) and with MCI/AD (Murty et al., 2021). Fifty-two subjects were retested with the same visual paradigm after a period of approximately one year. We employed large visual stationary Cartesian grating stimuli that induced two distinct gamma oscillations in both monkeys and humans (Murty et al., 2018), called slow [20-35 Hz] and fast gamma [36-66 Hz]. We estimated reliability of power in alpha and both gamma bands using a widely used measure called Intra Class Correlation (ICC; (Shrout and Fleiss, 1979)). We examined the test-retest reliability of SSVEPs at 32 Hz as well. We also studied correlations in the temporal and spectral profiles across same versus different individuals. In addition, we studied inter-subject variability (subject distinctiveness) through a linear classifier that was trained to discern same subject pairs from the all possible cross-subject pairs and compared the contribution of various features towards classification.

## Materials and Methods

### Human subjects

The full dataset was collected from 257 elderly human subjects (females: 106) aged 50 to 88 years as part of the Tata Longitudinal study of aging (TLSA). The subjects were recruited from urban Bengaluru communities, with careful evaluation by trained psychiatrists, psychologists and neurologists affiliated with National Institute of Mental Health and Neuroscience (NIMHANS) and M.S. Ramaiah Hospital, Bengaluru. Their cognitive performance was evaluated by a set of tests – ACE (Addenbrooke’s Cognitive Examination), CDR (Clinical Dementia Rating) and HMSE (Hindi Mental State Examination). More details have been provided in previous reports (Murty et al., 2020, 2021). Out of the 257 subjects recruited in the first year, we discarded data of 10 subjects due to noise. Out of the remaining 247 subjects, 52 subjects were retested after approximately a year. The test-retest sessions of a subject are referred to as baseline and follow-up.

All subjects took part against monetary compensation during both sessions. Informed consent was obtained from the subjects prior to the experiment. All procedures were approved by The Institute Human Ethics Committees of Indian Institute of Science, NIMHANS, Bengaluru and M.S. Ramaiah Hospital, Bengaluru.

### Experimental setting and behavioural task

Experimental setup and details of recordings have been explained in detail previously (Murty et al., 2020, 2021) and are briefly summarised here. EEG was recorded from 64-channel active electrodes (actiCap) using BrainAmp DC EEG acquisition system (Brain Products GMbH) and were placed according to the international 10-10 system. Raw signals were filtered online between 0.016Hz (first-order filter) and 1kHz (fifth order Butterworth filter), sampled at 2.5kHz, and digitized at 16-bit resolution (0.1µV/bit). It was subsequently decimated to 250 Hz. Average impedance of the final set of electrodes was (mean±SEM) 7.89 ± 0.67 kΩ and 7.73 ± 0.54 kΩ for baseline and follow-up, respectively. All the EEG signals recorded were referenced to electrode FCz during acquisition (unipolar reference scheme).

All subjects sat in a dark room facing a gamma-corrected LCD monitor (dimensions: 20.92 x 11.77 inches; resolution: 1289 x 720 pixels; refresh rate: 100 Hz; BenQ XL2411) with their head supported by a chin rest. It was placed at (mean± SD) 58 ± 0.7 cm from the subjects (range: 54.9 −61.0 cm) based on their comfort and subtended 52° x 30° of visual field for full screen gratings.

In the “gamma” experiment, subjects performed a visual fixation task, wherein 2-3 full screen grating stimuli were presented for 800ms with an inter-stimulus interval of 700ms after a brief fixation period of 1000ms in each trial using a customized software running on MAC OS. The stimuli were full contrast sinusoidal luminance achromatic gratings with either of the three spatial frequencies (1, 2, and 4 cycles per degree (cpd)) and four orientations (0°, 45°, 90°, and 135°). Out of the 52 subjects that were retested, 1 subject was not healthy (based on clinical scores), 3 subjects did not have a valid CDR score and 1 subject had unusable follow-up data, leaving with 47 pairs of subjects. With further constraints on the number of good electrodes for analysis (see Artifact Rejection) over both baseline and follow-up, we finally had 40 pairs of subjects (20 males and 20 females). Subjects with data available on both the sessions (N=40) aged in the range of (mean ± SD) 64.3 ± 7.17. The test-retest interval was (mean+SD) was 380.4±10.12 days (min: 256, max: 519), as reported in the plots in Figure 1 and Figure 2.

**Fig. 1.**
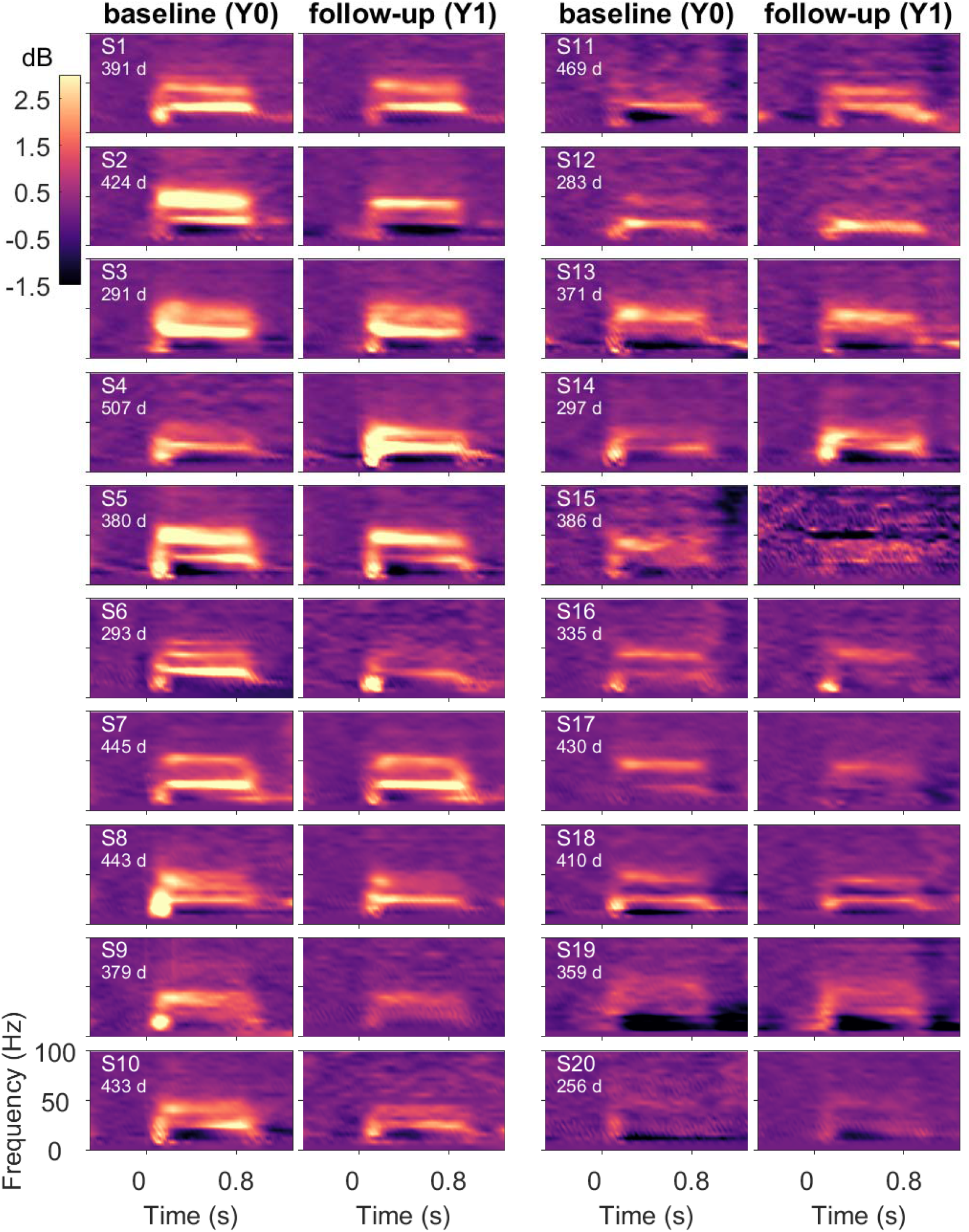
Gamma oscillations remain consistent across baseline and follow-up sessions. Change in time-frequency power spectra for the two recording sessions, baseline (left), and follow-up (right) of 20 female subjects to static gratings (“gamma” experiment). Subjects are ordered vertically based on the decreasing average slow and fast gamma power in visual electrodes, starting from the top left. The color bar on the top left denotes the log power ratio in dB units. The numbers indicate test-retest interval in days.

**Fig. 2.**
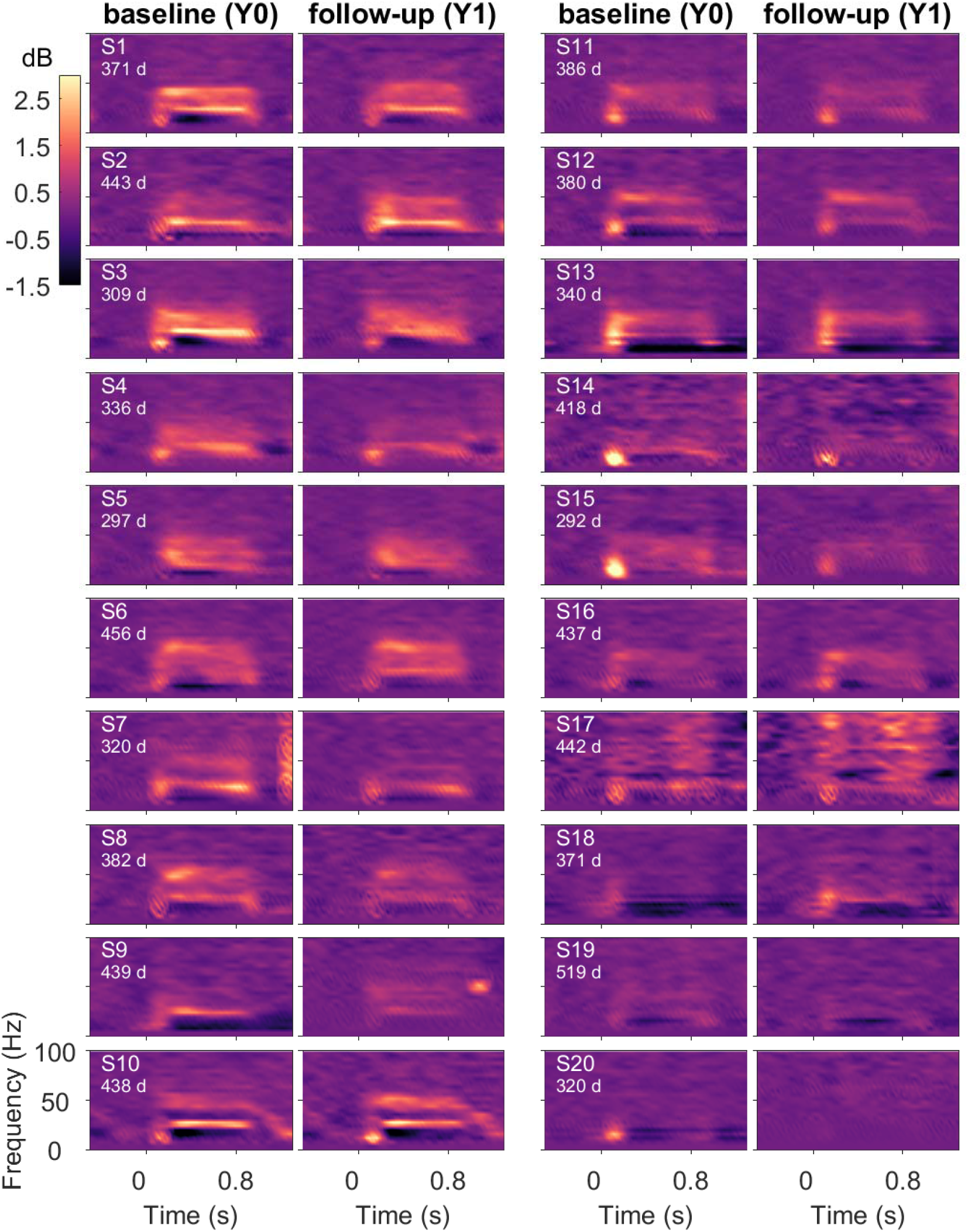
Same as Fig. 1 for 20 male subjects.

The “SSVEP” experiment was performed on a subset of these subjects. This was always conducted after the gamma experiment, on the same day. Here, subjects viewed a grating counter-phasing at 16 Hz, with the similar stimulus paradigm as gamma experiment, with spatial frequency and orientation that produced the maximum gamma in the gamma experiment. Out of the 52 subjects that were retested, 1 subject was not healthy (based on clinical scores), 3 subjects did not have a valid CDR score, 12 subjects either had unusable baseline or follow-up data, leaving us with 36 pairs of subjects (20 males and 16 females). Of these, pairs with number of trials less than 10 were discarded (4 subjects), ending up with 32 pairs (17 males and 15 females).

Eye position was monitored using EyeLink 1000 (SR Research Ltd), sampled at 500Hz. Any eye-blinks or change in eye position outside a 5° fixation window during −0.5s to 0.75s from stimulus onset were noted as fixation breaks, which were removed offline. This led to a rejection of 14.4±2.7% (mean±SEM) and 17.2±2.9% (mean±SEM) repeats for the baseline and the follow-up groups for the gamma experiment, and 14.9±2.7% and 16.5±3.0% for the SSVEP experiment.

### Artifact Rejection

We implemented a fully automated artifact rejection framework (for details, see (Murty et al., 2020, 2021), in which outliers were detected as repeats with deviation from the mean signal in either time or frequency domains by more than 6 standard deviations, and subsequently electrodes with too many (30%) outliers were discarded. Subsequently, stimulus repeats that were deemed bad in the visual electrodes (see the section on EEG data analysis below) or in more than 10% of the other electrodes were considered bad, eventually yielding a set of good electrodes and common bad repeats for each subject. Overall, this led us to reject (mean±SEM) 14.6±0.8% and 14.5±0.8% of repeats for the baseline and follow-up groups for the gamma experiment, and 6.8±0.7% and 7.1±0.8% for the SSVEP experiment. Finally, as in our previous studies, we computed slopes for the baseline power spectrum between 56-84Hz range for each unipolar electrode and rejected electrodes whose slopes were less than 0. We further discarded protocols which did not have any of the bipolar electrode pair (see EEG data analysis) in the visual electrode group after electrode rejection. The subjects with no good protocols (in either baseline or follow-up group or both) were rejected from analysis, yielding 40 subjects (20 females) for the gamma experiment and 32 subjects for the SSVEP experiment.

In addition to this artifact detection pipeline that was used in previous studies, two additional conditions were included. First, for each subject, we computed the union of bad electrodes in baseline and follow-up sessions and rejected this union from both sessions, such that analyses for baseline and follow-up were over the same set of electrodes. Second, we noticed the presence of a large discontinuity in the signal in a small fraction (∼3%) of stimulus repeats and added a pipeline to remove those. Specifically, stimulus repeats with discontinuities (in any of the electrodes) were detected by thresholding the derivative of signal (> 120 μV/sec) in time within [-0.5 1.5] sec with respect to stimulus onset. This led to the rejection of (mean±SEM) 5.07±0.89 (maximum: 21) repeats in baseline and 4.55±1.04 (maximum: 23) repeats in follow-up subjects. Electrodes with more than 20 such discontinuities were considered noisy (maximum: 5 electrodes). These additional conditions were added mainly to improve the time-frequency plots (Figures 1-2) that were sensitive to these large discontinuities; the main results (Figures 5-8) remained similar even without these additional conditions. Overall, after discarding all bad repeats, 147.3± 6.0 and 152.8±6.4 stimulus repeats were available for the gamma experiment, and 63.2± 2.0 and 61.1±2.5 for the SSVEP experiment, for the baseline and the follow-up groups, respectively.

### EEG data analysis

All analyses were performed using bipolar reference scheme. Every electrode was re-referenced offline to its neighbour, yielding 112 bipolar pairs of electrodes from the 64 unipolar electrodes (Murty et al., 2020, 2021). The bipolar pairs among the visual electrodes considered for analysis (except for scalp maps) were: PO3-P1, PO3-P3, POz-PO3 (left anterolateral group); PO4-P2, PO4-P4, POz-PO4 (right anterolateral group) and Oz-POz, Oz-O1, Oz-O2 (posteromedial group).

All the data analyses were done using custom codes written in MATLAB (MathWorks. Inc; RRID:SCR_001622). Power spectrum and time-frequency spectrograms were obtained using the Chronux Toolbox ((Bokil et al., 2010), RRID:SCR_005547). For analysis, similar to our previous study (Murty et al., 2020), we chose [-500 0] msec as the baseline period and [250 750] msec as the stimulus period, yielding a resolution of 2Hz.. This stimulus period was so chosen to avoid stimulus-onset transients. Time-frequency spectrogram was computed with 4Hz frequency resolution (moving window of 250ms) and a step size of 25ms.

Change in power between stimulus period and baseline period for a frequency band was computed using the following formula:

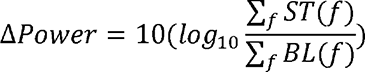

where ST is the stimulus power spectra and BL is the baseline power spectra, both averagedithin relevant frequency bins (*f*), across all the analyzable repeats. Band powers are specifically computed in three frequency bands, namely slow gamma (20-35 Hz), fast gamma (36-66 Hz), and alpha (8-12 Hz). These were averaged across all the three electrode groups. Scalp maps were generated using the *topoplot* function of EEGLAB toolbox ((Delorme and Makeig, 2004), RRID:SCR_007292) with custom bipolar montage of 112 channels.

Test-retest reliability was assessed by Intra-Class Coefficient (ICC) measure. Suppose the dataset is arranged in a matrix with each column representing a session and each row a different subject. ICC incorporates both inter-subject and intra-subject variability in terms of mean square (variance) for rows (MSR) [variance of sessions’ means across subjects] and mean square (variance) within raters (MSW) [mean of subjects’ variance across sessions] respectively. A specific type of ICC, called 1-1 ICC, which is based on one-way analysis of variance model, is defined as follows (see (Shrout and Fleiss, 1979) for details):

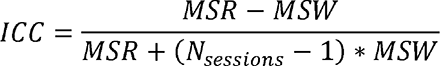

In our case, since = 2, this reduces to ICC = (MSR-MSW)/(MSR+MSW). This measure increases with decreasing within-subject variability (MSW) and with increasing between-subject variability (MSR), with values approaching 1 with perfect reliability (MSW=0). This measure was used for the reliability of spectral power in sensor and source domains in MEG (Martín-Buro et al., 2016). Also, the null-hypothesis : H_O_: ICC=0 was tested for significance by *F*-statistical test, *F-*value = MSR/MSW, with (*N_subjects_* −1) and *N_subjects_* (*N_sessions −1_*) degrees of freedom (Shrout and Fleiss, 1979), reported with 95% confidence interval, F-value and p-value. Larger the variation of session means across subjects, relative to the variation within sessions, larger is the *F*-value. Given the *F*-value, degrees of freedom of MSR and MSW, p-value is computed based on the *F* cumulative distribution function. Two related types of ICC, called Consistency and Agreement ICC (ICC(C,1) and ICC(A,1)) were also computed for comparing our results with other studies. These ICC definitions involve MSE (Mean Square for Error) which encompasses variance across both subjects and sessions (for details, see (McGraw and Wong, 1996)). We computed the intraclass correlation using MatlabCentral file-exchange codes provided by Salarian (2016), which implemented the statistical testing described by (McGraw and Wong, 1996).

### Estimating subject distinctiveness using linear discriminant analysis (LDA)

A measure of subject distinctiveness reflects the separability between the self-pairs and cross-pairs—which is approximated here, in terms of classifier performance (Linear Discriminant classifier) in discriminating the two categories of pairs. Input to the classifier is a pair of baseline and follow-up session recordings from two subjects. For each pair, we defined seven comparative measures (stimulus induced features), namely, absolute difference in relative band powers within slow gamma (p_sγ_) [20-35 Hz], fast gamma (p_fγ_) [36-66 Hz], and alpha (p_α_) [8-12 Hz] (3 features), baseline corrected band power temporal correlations (Pearson’s correlation) in the slow gamma (tc_sγ_), fast gamma (tc_fγ_), and alpha (tc_α_) between [0 0.8] secs (3 features; correlation between the traces shown in Figure 3), and spectral correlation (*sc*) in [0-100Hz] (excluding 50 Hz and 100 Hz peaks that represented line noise and monitor refresh rate; 1 feature; correlation between the traces shown in Figure 4). To study the importance of the features in the baseline period, we also considered baseline alpha difference power (p_bl__α_) and raw PSD correlation (*bl sc*) features (correlation between the baseline PSDs shown in Supplementary Figure 3). Three binary classifiers (using only stimulus-induced features, only baseline features, and both feature sets combined) were trained to classify a pair into either self-pair or cross-pair and was tested according to 5-fold stratified cross-validation.

**Fig. 3.**
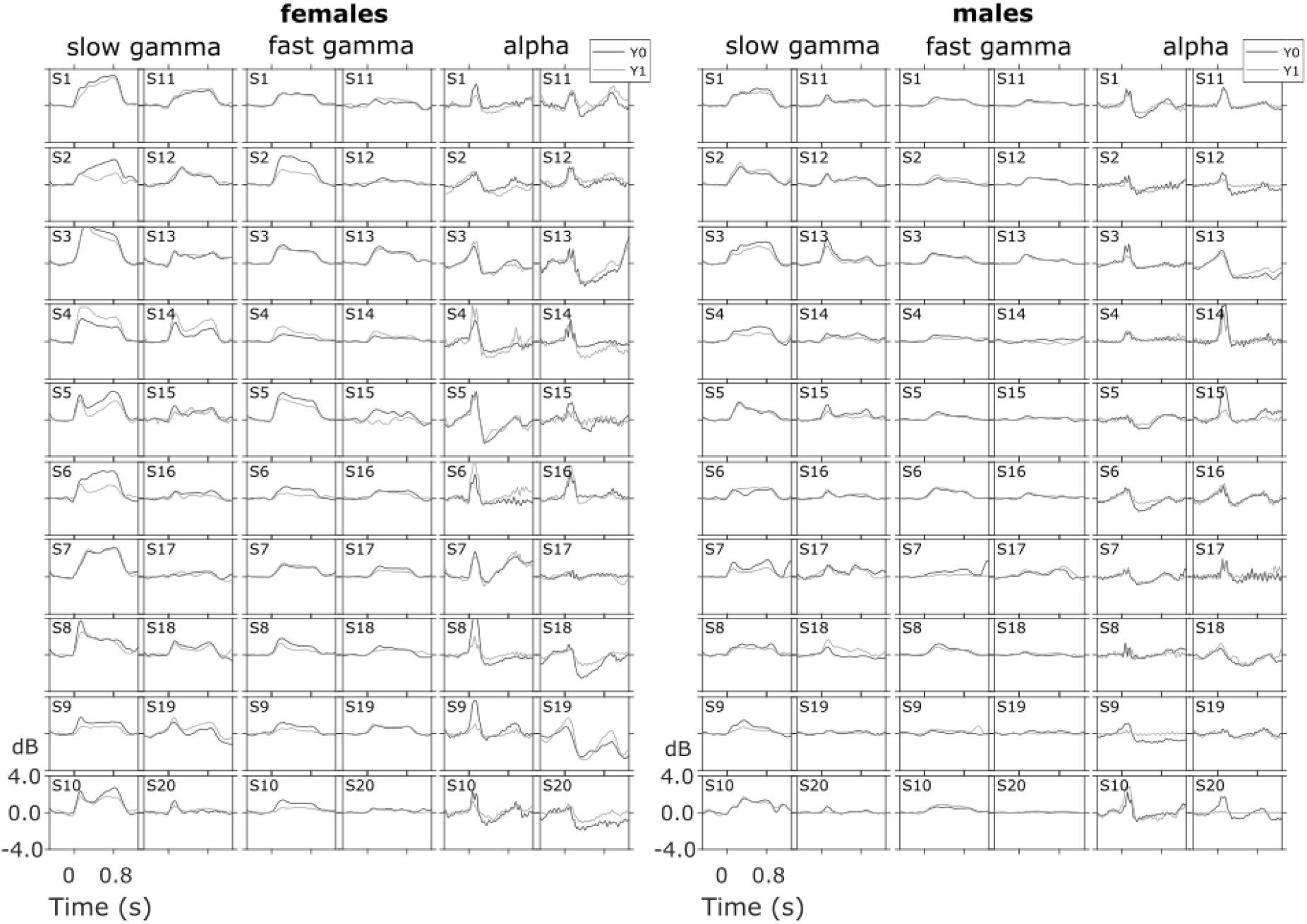
Temporal evolution of band powers in slow gamma, fast gamma and alpha range across baseline and follow-up remain consistent in males and females. Band power as a function of time, computed from averaging baseline corrected time-frequency spectra across relevant frequency bins for slow gamma, fast gamma and alpha bands. Traces are plotted in black for baseline (Y0) and gray for follow-up (Y1) for 20 female and 20 male subjects. Same subject order as in previous plots.

**Fig. 4.**
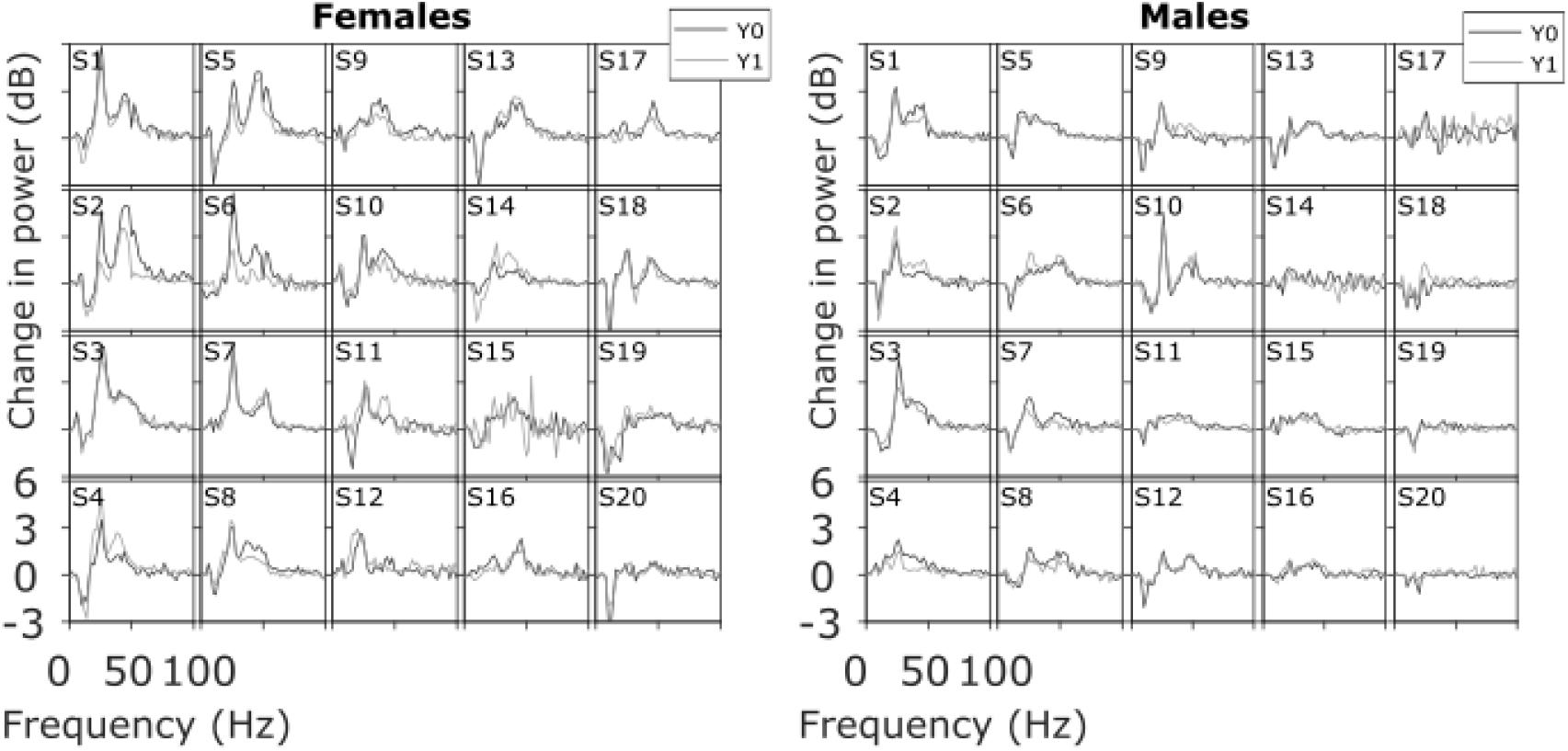
Power spectral densities (PSDs) are consistent across baseline and follow-up in males and females. The change in PSD during stimulus period relative to baseline PSD for baseline (Y0; black) and follow-up (Y1; gray) for 20 female and 20 male subjects.

Because the cross-pairs (40×39 = 1560) far outnumbered the self-pairs (40), classifier performance could not be assessed using accuracy, since even a null classifier that categorizes every observation as a cross-pair would have an accuracy of 97.5%. Instead, we assessed the classifier performance through the area under the ROC (Receiver Operating Characteristic) curve (AUC), where the ROC curve represents the true positive rate against false positive rate. AUC calculation does not need the class label output of classifier, as it considers various thresholds for classification. Only the weighted projection along with the true class labels are needed for true positive rate and false positive rate. AUC values vary between 0.5 (chance level classification) and 1 (perfect classification). Feature importance was determined by the weight of the feature computed using LDA classifier on the z-scored feature values, as well as using single feature AUC method wherein each feature was individually considered to classify self versus cross pair. We used MATLAB statistics and machine learning toolbox function *perfcurve.m* for computing AUC.

### Statistical Analysis

All the statistical analysis and results obtained, are using the power spectrum or correlation measures. Appropriate tests (t-statistic; F -statistic; Wilcoxon rank sum) were used to interpret our findings.

### Data and code availability

All spectral analyses were performed using Chronux toolbox (version 2.10), available at http://chronux.org. Intraclass correlation analysis functions were taken from MatlabCentral file-exchange, provided by Salarian (2016). Data used here was made available in our previous study (Murty et al., 2021) and the codes would be made available to readers upon request and publicly available at a later time.

## Results

### Time frequency spectra remain reliable over one-year interval

We first examined the change in time-frequency power spectra relative to baseline power (−0.5 to 0 seconds) of EEG signals for the “gamma” experiment, averaged across the three bipolar electrode groups, for the baseline and follow-up sessions (see Methods). Figures 1 and 2 show the results for female (N=20) and male subjects (N=20), sorted based on decreasing total gamma power. Both females and males show visually reliable time-frequency spectra within a subject across baseline (left panel) and follow-up (right panel). Specifically, the powers in the three frequency bands, slow gamma, fast gamma, and alpha were consistent in time-frequency spectra, as were the temporal evolution of these rhythms. Similarly, the topographic scalp maps of average gamma band power (mean of slow and fast gamma) appeared consistent between baseline and follow-up sessions in females (Supp. Fig. 1) and males (Supp. Fig. 2).

### Band powers, spectral and temporal profiles all remain highly reliable

Figure 3 shows the temporal evolution of band powers in alpha, slow gamma, fast gamma range, computed by averaging time frequency spectra across relevant frequency bins, separately for males and females. Remarkably, these profiles were distinct across subjects but appeared highly correlated across the two sessions within subjects. Figure 4 shows the change in power during the stimulus period (0.25 – 0.75 seconds, where 0 is the stimulus onset) relative to baseline for the two sessions. These traces show a prominent suppression at alpha range, and elevation of power at slow and fast gamma bands, and together represent a “spectral profile” for each subject. These spectral profiles were also highly similar for each subject and distinct across subjects. Consistency was found in the spectral profile of baseline period absolute power as well, although for several subjects there was substantial difference in the noise floor (Supp. Fig. 3). For these subjects, the stimulus-induced PSDs were also different in a similar way (data not shown), such that the change in PSD plots shown in Figure 3 remained highly overlapping. Further, in spite of the difference in noise floor, the spectral shape during baseline periods was informative about subject identity (see Figure 8 for details).

Figure 5a-c show a scatter plot of power in slow gamma, fast gamma and alpha bands in baseline versus follow-up sessions. Supp. Fig. 4 shows similar scatter plot of average resting state alpha power (baseline period). SSVEP power (increase in power at 32 Hz from baseline) computed from the SSVEP experiment (see Methods) is also shown (Fig. 5d). In all cases, the powers were highly correlated (significance results are indicated in the plots).

**Fig. 5.**
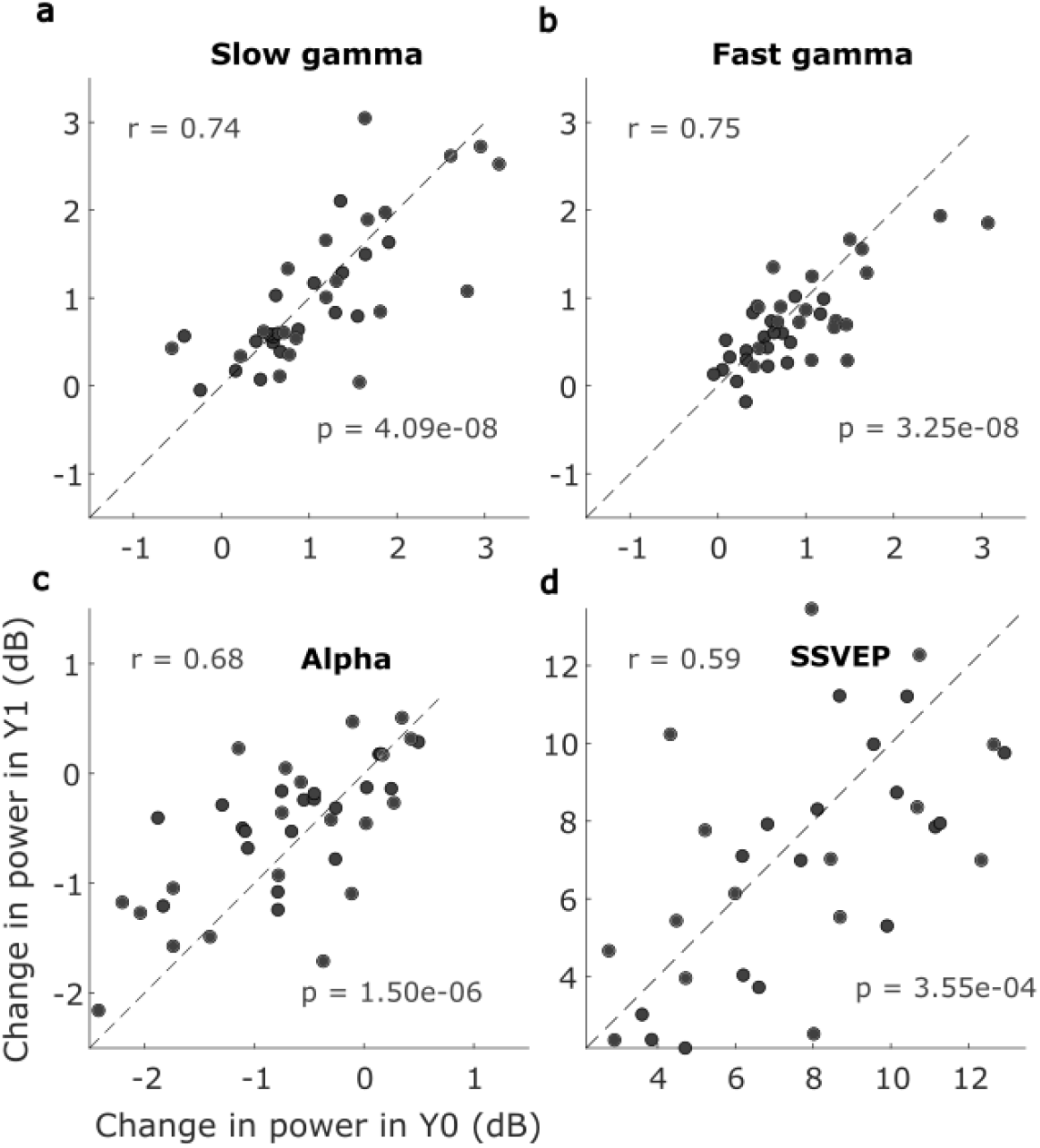
Change in slow gamma, fast gamma, alpha power and SSVEP at 32 Hz power are correlated across baseline and follow-up. (a) – (c) Scatter plots of the average change in power within slow gamma, fast gamma and alpha bands for baseline and follow-up sessions. (d) The average change in power at SSVEP frequency at 32 Hz. Pearson’s correlation coefficient and p-value are mentioned on the top left and bottom right in each panel. Note the difference in axis limits across the plots.

We further assessed the reliability using a metric called ICC (Shrout and Fleiss, 1979), which compares the ratio of variances within and across subjects. Average band powers were highly reliable for all frequencies ranges: slow gamma ICC: 0.74 (95% CI = 0.56–0.85; F = 6.77, p = 9.68e-9), fast gamma ICC: 0.69 (95% CI = 0.50–0.82; F = 5.64, p =1.41e-7), alpha ICC: 0.65 (95% CI = 0.43–0.79; F = 4.68, p =1.89e-6) and SSVEP ICC: 0.58 (95% CI = 0.30–0.77; F = 3.75, p =1.76e-4). In addition, average resting state alpha power was also reliable; ICC: 0.78 (95% CI = 0.62-0.88; F = 8.12, p = 5.94e-10). Previous studies have used two other variants of ICC, called consistency ICC(C,1) and agreement ICC(A,1), see Methods and (McGraw and Wong, 1996) for details. These measures were also comparable to the classical ICC in our dataset. For slow gamma, fast gamma, alpha and SSVEP, ICC(C,1) values were 0.74, 0.72, 0.67, and 0.59, respectively, while ICC(A,1) values were 0.74, 0.70, 0.65, and 0.58.

To quantitatively assess the reliability in the temporal evolution and spectral profile, we computed the Pearson’s correlation between baseline and follow-up traces for each subject (“self-pair correlation”) and compared with the average correlation between the follow-up traces of all other subjects (“cross-pair correlation”). For the temporal correlation of slow gamma, self-pair correlation was 0.87±0.09 (median±median absolute deviation (MAD)), significantly greater than cross-pair correlation (0.28±0.09; Left-sided Wilcoxon rank sum test, p=1.59e-13, N=40/1560) (Fig. 6a). Similarly, for the temporal correlation of fast gamma, self-pair correlation was 0.88±0.09 (median±MAD), significantly greater than cross-pair correlation (0.73±0.07; Left-sided Wilcoxon rank sum test, p=8.74e-7, N=40/1560) (Fig. 6b). Likewise, alpha temporal correlation within self-pair was 0.94±0.03 (median±MAD), significantly greater than cross-pair correlation (0.88±0.03; Left-sided Wilcoxon rank sum test, p=2.1e-7, N=40/1560) (Fig. 6c). For spectral correlation, self-pair correlation was 0.85±0.08 (median±MAD), significantly greater than cross-pair correlation (0.56±0.08; Left-sided Wilcoxon rank sum test, p=3.36e-13, N=40/1560) (Fig. 6d).

**Fig. 6.**
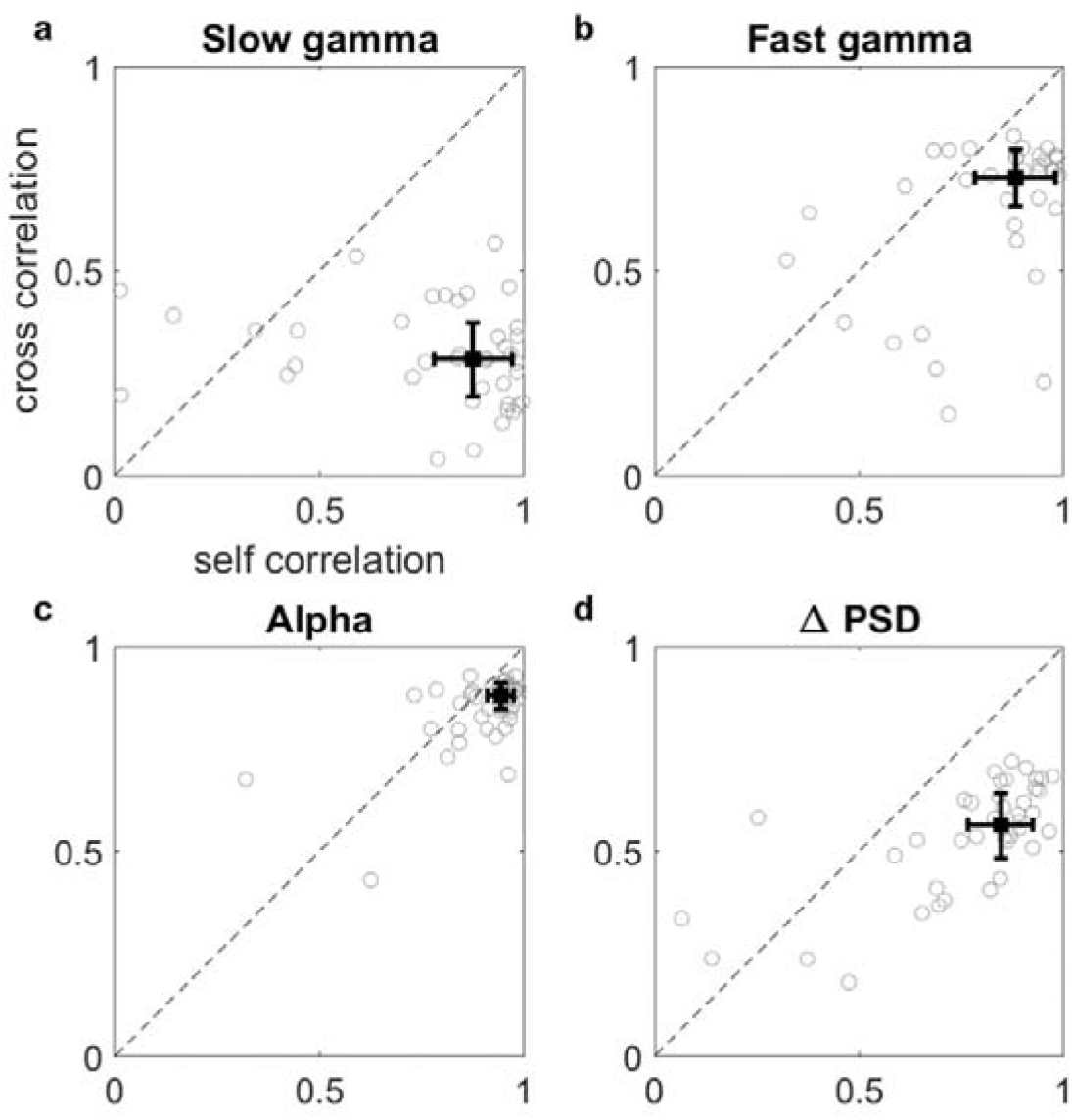
Change in PSD profile and temporal profiles of band powers are more correlated within self-pairs than cross-pairs. (a) – (c) Correlation between the band power time series traces (shown in Figure 3) computed between two sessions of the same subject (self-pair) plotted against the median correlation between traces for all other subjects (cross-pair), for slow gamma (a), fast gamma (b) and alpha (c) bands. (d) Same plot for the change in PSD traces shown in Figure 4. There are 40 data points in each plot. The median of these data is shown by the black box with error bars indicating median absolute deviation (MAD).

### Feature based subject identification

From the data collected under “gamma” experiment, we computed subject distinctness measures, reflecting the separation between the self-pairs (40/1600) and cross-pairs (1560/1600), using spectral and temporal profiles of the signal. The separation was assessed within a classifier framework, based on seven stimulus-induced features including band power differences (slow gamma, fast gamma and alpha bands; 3 features), temporal profiles (3 features), and spectral profile (1 feature; Fig. 7; see Methods for details) using Linear Discriminant Analysis (LDA). Feature weights were computed by 5-fold stratified training and was used for classifying test data.

**Fig. 7.**
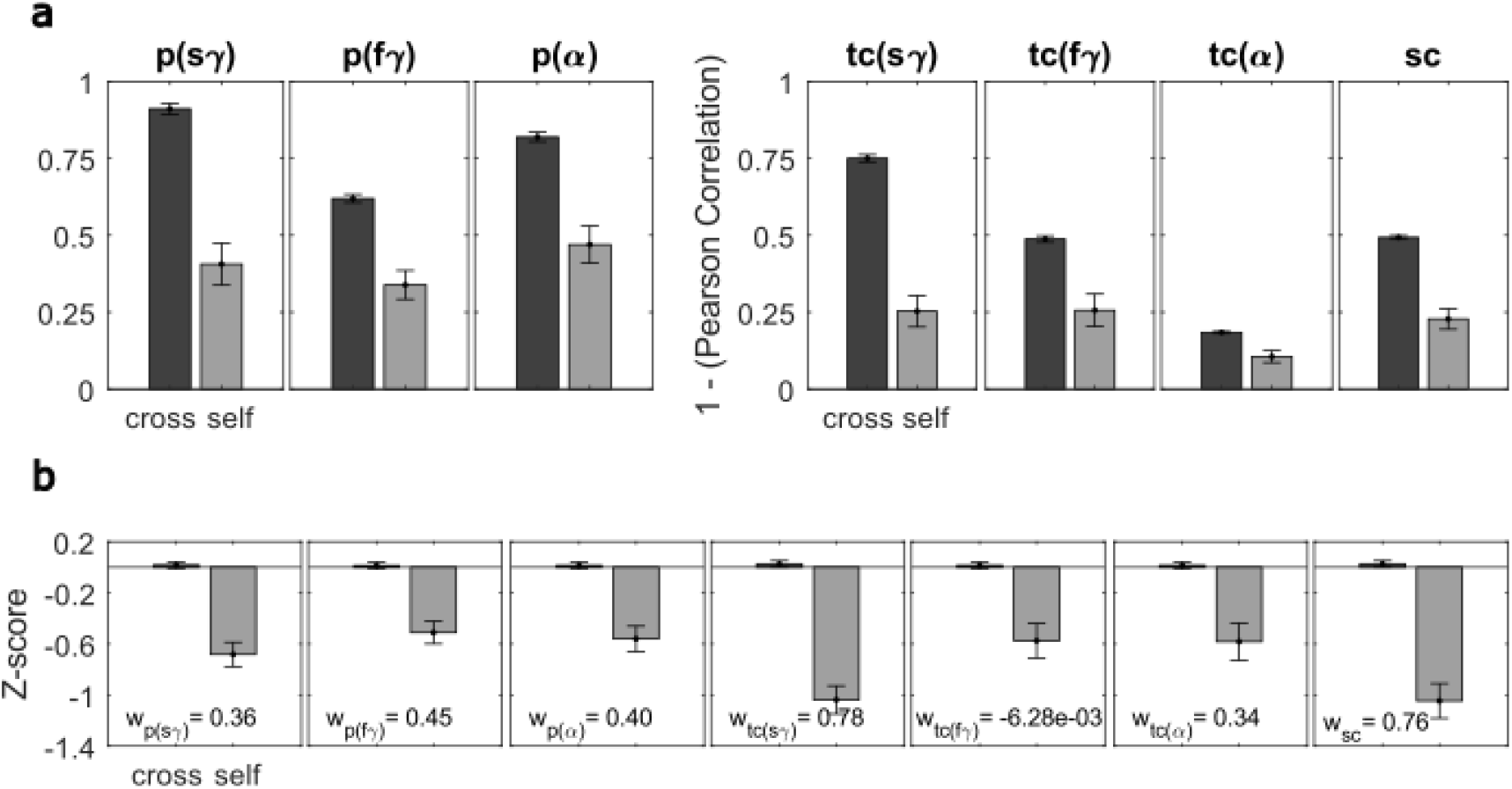
Temporal slow gamma and spectral correlations emerge as dominant features in classification. (a) The seven features provided as inputs for the subject distinctness analysis are plotted separately for cross-pairs (cross individual comparisons) and self-pairs (same individual comparisons). Change in power (first 3 features) are in units of dB. Temporal and spectral correlations (other four features) are unitless. (b) Z-scored features are plotted in the same way as panel (a). The values mentioned in the figure insets denote the classifier weight of the respective feature in separating out self-pairs from the cross-pairs.

Figure 7a shows the mean values of the seven features for the cross and self-pairs. As expected, the band power differences were lower for self-pairs than cross-pairs (plots on the left). To use the correlation based features in the same way to reflect dissimilarity, we subtracted 1 from the correlation values so that they were also lower for self-pairs than cross-pairs (right plots). To compare the distinctness of these features, and for data standardization, we z-scored each feature value across all the 1600 (40×40) pairs (Figure 7b). All features had lower z-scores for self-pairs compared to cross-pairs, with the temporal profile of slow-gamma and the spectral profile having the largest separation in z-scores.

Next, we performed LDA on these z-scored data. Because data are z-scored, the weights of the classifier (shown in the legends of Figure 7b) can be compared directly and reflect the importance of the features towards classification. These weights were also the highest for the temporal profile of slow-gamma and the spectral profile. Note that these weights are not always proportional to the difference in z-scores between the two classes, since some features may be highly correlated with others and may therefore carry redundant information (for example, temporal variation in fast gamma had a weight of almost zero).

Further, we assessed the performance of each feature separately by calculating the AUC, which is a measure of the separation between the data in the two classes that does not depend on any explicit threshold (Figure 8). Again, AUC values were highest for the temporal correlation of slow-gamma (0.84) and spectral correlation (0.83). However, including all seven features improved the AUC only marginally to 0.89, suggesting that these features had high redundancy. We also considered the baseline features (features obtained from resting state) including absolute average alpha power (1 feature; from the PSD in baseline period, Supp. Fig. 3) and spectral profile (1 feature; correlation between the PSDs shown in Supp. Fig. 3; see Methods for details) to determine the improvement in classification with inclusion of stimulus-induced features. Features obtained from resting state data also performed well (AUC=0.82), with resting state alpha power (AUC=0.73) performing better than spectral correlation of baseline PSDs (AUC=0.67). The features in the baseline period were redundant with the stimulus features; adding these baseline features only improved the AUC to 0.90 (Figure 8). These observations establish slow gamma temporal correlation and spectral correlation as the dominant features.

**Fig. 8.**
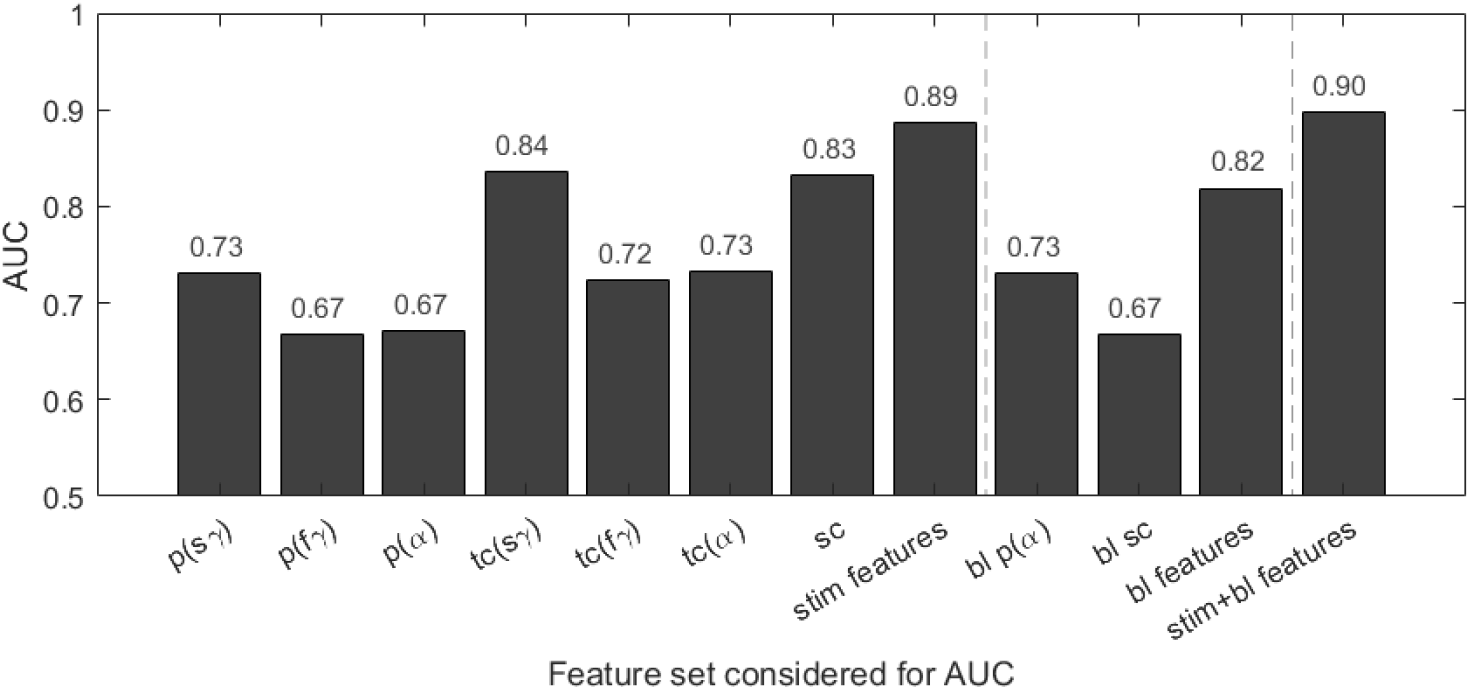
Features from stimulus period perform better than resting state features. The AUC values obtained using various classifiers, constructed based on the feature set listed on the x-axis. First, the performance of each of the stimulus-induced features is considered individually and subsequently collectively, and then the baseline (resting state) features are considered, first individually and subsequently collectively, and finally combined with the stimulus induced features.

## Discussion

We studied reliability of various spectral and temporal neural markers on visual stimulus induced responses in EEG recorded from elderly (>49 years) healthy human subjects, with a test-retest interval of about one year. Time frequency spectra were consistent across baseline and follow-up sessions for both males and females. Similar consistency was observed for temporal traces of alpha, gamma band powers as well as change in PSD profiles. We used intraclass correlation (ICC) to measure reliability and found power in alpha, slow and fast gamma to be significantly reliable. SSVEP power at 32Hz were also found to be reliable. Temporal and spectral profiles were tested for reliability through Pearson’s correlation between the profiles of same individual pairs (“self-pair correlation”) and were found to be significantly higher than the correlations between cross individual pairs (“cross-pair correlation”). We trained a linear classifier to distinguish pairs of datasets as belonging to the same subject (self) versus different (cross) using seven features including absolute difference of powers (3 features), temporal band power correlations (3) and spectral correlation (1). The feature weights as well as the separation in individual feature z-scores were highest for slow gamma temporal correlation and spectral correlation. We measured classifier performance using area under ROC curve and found that each of these two features also performed the best (AUC of ∼0.84). Overall classification performance increased only modestly when all features were used (AUC of 0.89), suggesting high redundancy between features. Classification using only resting state (baseline) features was also high (AUC of 0.82 when resting alpha and baseline PSDs were considered) and combining resting state and stimulus-induced features yielded AUC of 0.90. Overall, our results suggest that stimulus induced spectral and temporal changes in various frequency bands are reliable and distinct, allowing their use in subject identification as well.

### Previous studies

#### Reliability of fast and slow gamma

Visually induced gamma oscillations in fast gamma range were shown to be reliable in human MEG with substantially higher ICC(C,1) of 0.89 (Hoogenboom et al., 2006; Tan et al., 2016) compared to our results (ICC(C,1) of 0.72 for fast gamma). However, these two studies had much lower test-retest period of 1-11 day (Tan et al., 2016) and 4 days to 4 months (Hoogenboom et al., 2006), and the stimulus was a contracting circular sine wave grating. Hence, the low ICC in our case could be due to multiple reasons, such as the stimulus, recording modality and the test-retest period. Test-retest interval seems to be an important factor, since McCusker and colleagues (2021) who studied induced gamma in MEG after a period of up to 3 years reported much lower agreement ICC(A,1) of 0.55 (McCusker et al., 2021). Apart from the duration, the lower values could be due to the ICC type used, as the agreement type ICCs are usually lower than the consistency and classical definitions in presence of systematic bias in the data (although we did not find this in our data). Moreover, it could be due to the use of a checkerboard stimulus. Another human MEG study showed test-retest reliability in induced gamma oscillations with one week gap (Muthukumaraswamy et al., 2010), but they did not report ICC values.

This is the first time that the reliability of induced slow gamma oscillations was addressed in human EEG with a gap of approximately a year. Slow gamma was observable in a single subject (S02) in the study by Tan and colleagues (2016) in human MEG, but they do not follow up on it further. They used many stimuli with differences in spatial structure (circular or Cartesian grating), temporal structure (moving or stationary), and size (large or small), however, only some of which produced slow gamma. We have previously shown that slow gamma oscillations are induced in human EEG with stationary Cartesian gratings of large size (Murty et al., 2018) and hence we employed similar stimulus parameters for this study.

#### Reliability of alpha band

Corsi-Cabrera and colleagues (2007) found less inter-session reliability in absolute alpha band power with eyes open compared to beta band. They attributed less inter-session reliability to the higher reactivity of alpha to external stimulation modifying the subjects’ internal state. Several studies have compared absolute versus relative power (percentage value of power within the band w.r.t total power across all the bands), with mixed results. Some studies found relative alpha power to be more reliable than absolute power (John et al., 1983; Kondacs and Szabó, 1999), others found comparable reliability (Gasser et al., 1985; Salinsky et al., 1991), while a recent study (McCusker et al., 2021) reported more robust reliability of absolute alpha activity in resting state than in the stimulus period (alpha desynchronization). In our study, relative power (ICC(1) = 0.81) measures were more reliable than absolute power (ICC(1) = 0.76) in the alpha band, within the stimulus period. Moreover, in line with the study by McCusker and colleagues (2021), the resting state alpha reliability of both the relative power (ICC(1) = 0.83) and absolute power (ICC(1) = 0.78) were higher than the stimulus period. Apart from alpha power, alpha peak frequency also has been investigated and found to be test-retest reliable (Maltez et al., 2004; Näpflin et al., 2007, 2008; Grandy et al., 2013).

#### Reliability of other signals

Test-retest reliability has been investigated in signals such as event-related synchronization and desynchronization (ERS/ERD) (Neuper et al., 2005), theta and alpha oscillations during a working memory task (McEvoy et al., 2000), functional EEG network characteristics (Velde et al., 2019) and evoked oscillations with visual stimuli (Keil et al., 2003; Fründ et al., 2007). Although the degree of reliability is difficult to compare across studies, all measures have shown some degree of test-retest reliability. Similarly, spectral features like the spectral exponent (slope of the PSD in log-log coordinates) have been shown to convey test-retest reliability (Demuru and Fraschini, 2020).

Although, to our knowledge, reliability of SSVEPs in gamma range has not been studied, there are several studies on its auditory equivalent, the auditory steady-state response (ASSR) (McFadden et al., 2014; Legget et al., 2017; Hirano et al., 2020). Interestingly, ASSRs have been shown to be abnormal in patients with autism (Wilson et al., 2007) and schizophrenia (Brenner et al., 2003). Hence, the aforementioned reliability studies have validated the use of ASSRs as a neuropsychiatric biomarker, including those employing MEG recordings (Tan et al., 2015). A recent study further showed ASSRs to be reliable in schizophrenia patients (Roach et al., 2019).

#### Subject distinctness

Näpflin and colleagues (2007) used alpha power, centre frequency, and spectral correlation (in [2-32 Hz]) for data collected under eyes closed and eyes open paradigm to separate out same individual pair from inter-individual pair using a Generalized Linear Model (GLM) and reported 88% sensitivity or probability of detection with 40 subjects. In their study, GLM was fitted for each person separately. We instead formulated a classifier over all the subjects and measured performance using AUC. They also reported remarkable performance by the spectral shape feature. Kondacs and Szabo (1999) found lower intra-individual than inter-individual variability mainly in alpha power (7.5-12.5 Hz) and alpha mean frequency in eyes closed condition. We chose spectral shape feature (spectral correlation) (Figure 7a) and alpha power (Supp. Fig. 4). Computation of alpha mean frequency was not informative in our case since the spectral resolution was 2 Hz (due to analysis duration of 0.5 seconds), giving rise to only 3 bins in alpha range (8-12 Hz). Hence, we did not consider alpha mean frequency feature in this study.

### Dominance of spectral profile and slow gamma temporal correlation and in classification

High performance of spectral profile feature is not unexpected, since this feature captures the changes in power at all frequencies including alpha and gamma bands, as well as features such as the shape of the PSD, and has been previously shown to perform well in classification tasks (Napflin et al., 2007). More surprising was the high performance of the temporal profile of slow gamma. As shown in Figure 3, first column, slow gamma temporal profile was more variable across subjects than other measures such as fast gamma. We have previously shown that unlike fast gamma which is induced soon after stimulus onset and then is subsequently maintained throughout the stimulus duration, slow gamma builds up slowly over time (Murty et al., 2018). On the other hand, the transient broadband component is high early after stimulus onset (Figures 1 and 2). Depending on the ratio of transient and steady-state components in the temporal profile, slow gamma temporal profile showed higher variability across subjects than fast gamma, which generally was highest after stimulus onset and then decayed (Figure 3).

Another reason could be the nature of slow gamma itself. Slow gamma has larger functional spread, i.e., larger coherent neural population in comparison with fast gamma, as shown in monkey LFP (Murty et al., 2018). Such large-scale synchrony might contribute to its relatively higher reliability. Indeed, slow gamma reduced more slowly with age compared to fast gamma (slope of band power versus age plots were shallower; (Murty et al., 2020)). High distinctness could arise from the larger subspace for variability across cortical space within the source network. In general, slow oscillations (in alpha and lower beta bands) have been related to larger information integration, whereas gamma oscillations have been associated with more local computations (von Stein and Sarnthein, 2000). A study in mouse V1 reported “spatial context-dependent” induced gamma rhythm at ∼30 Hz depending on the grating size (Veit et al., 2017). This rhythm was shown to be mediated by long-range somatostatin (SOM) interneurons, which are commonly associated with a multitude of neuropsychiatric diseases (Lin and Sibille, 2013). This highlights the necessity of checking slow gamma reliability for its usage as a biomarker for diagnosing neuropsychiatric diseases.

The dominance of slow-gamma over fast-gamma also has important practical considerations. In EEG, higher frequencies (>30 Hz) often get corrupted by a high noise floor (as observed in a few cases in our data as well; see Supp. Fig. 3). In addition, the line noise at 50 or 60 Hz is within the fast gamma range. These factors impede accurate estimation of fast gamma in EEG (which could have contributed to a lower performance in our data as well). Slow gamma does not suffer from these practical limitations. Therefore, using static, full field Gratings that induce strong slow gamma could be a convenient way to obtain a richer spectral signature that is also more reliable with age than fast gamma (Murty et al., 2020) and is equally compromised in Alzheimer’s Disease as fast gamma (Murty et al., 2021).

### Factors that influence reliability

Parvalbumin (PV) positive interneurons are thought to play a key role in the generation of fast gamma oscillations (Bartos et al., 2007; Cardin et al., 2009; Sohal et al., 2009). On the other hand, optogenetic stimulation of SOM interneurons induce slow gamma oscillations under some contexts in rodents, suggesting their role in generation of slow gamma. The stability of these inhibitory networks over time could determine the reliability of gamma oscillations. In addition, oscillations recorded on the scalp could be influenced by various subject dependent parameters like head geometry, scalp conductivity, and other low-level factors. Further, some studies have shown cortical thickness to be correlated with gamma peak frequency, but not with power values (Muthukumaraswamy et al., 2010; Gaetz et al., 2012). Auditory steady-state activity (ASSR) at 40 Hz was also shown to be positively associated with left superior temporal gyrus cortical thickness in healthy controls (Edgar et al., 2014). Since these external factors may also change with time, the reliability of the recorded signal reflects both the stability of the sources and these confounding factors. Some of these factors such as cortical folding (Schultz et al., 2013), cortical thickness (Gaetz et al., 2012), and surface area (Herting et al., 2015) can be estimated using subject specific magnetic resonance imaging (MRI). Moreover, reliability estimates could be improved by increasing the signal to noise ratio by using accurate head models allowing us to go from the sensor to source space (Hoogenboom et al., 2006; Muthukumaraswamy et al., 2010; Martín-Buro et al., 2016; Tan et al., 2016). Such improvements, as well as more refined techniques to capture network properties that go beyond power-based measures (Höller et al., 2017; Kuntzelman and Miskovic, 2017), hold promise towards a more advanced and reliable biomarker of mental disorders.

## Funding disclosure and Acknowledgements

This work was supported by Tata Trusts Grant, Wellcome Trust/DBT India Alliance (Senior fellowship IA/S/18/2/504003 to SR), and DBT-IISc Partnership Programme.

---

**Supp. Fig. 1.**
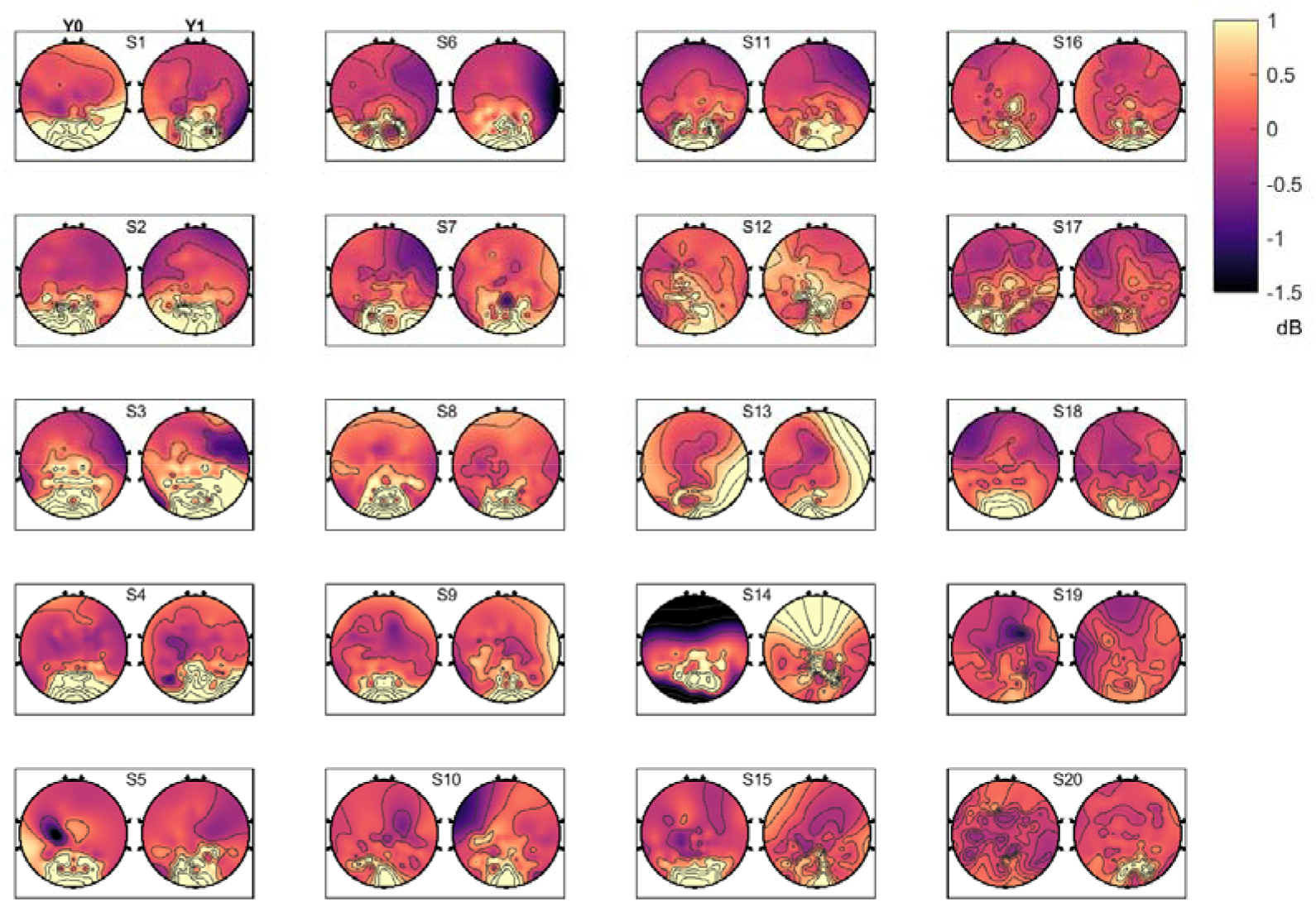
Scalp maps for average gamma power are also consistent between baseline and follow-up sessions. Average of slow and fast gamma power across common good bipolar electrodes is plotted as a scalp map for 20 female subjects. Same subject order as Figure 1. The color bar on the right denotes the power ratio in dB units.

**Supp. Fig. 2.**
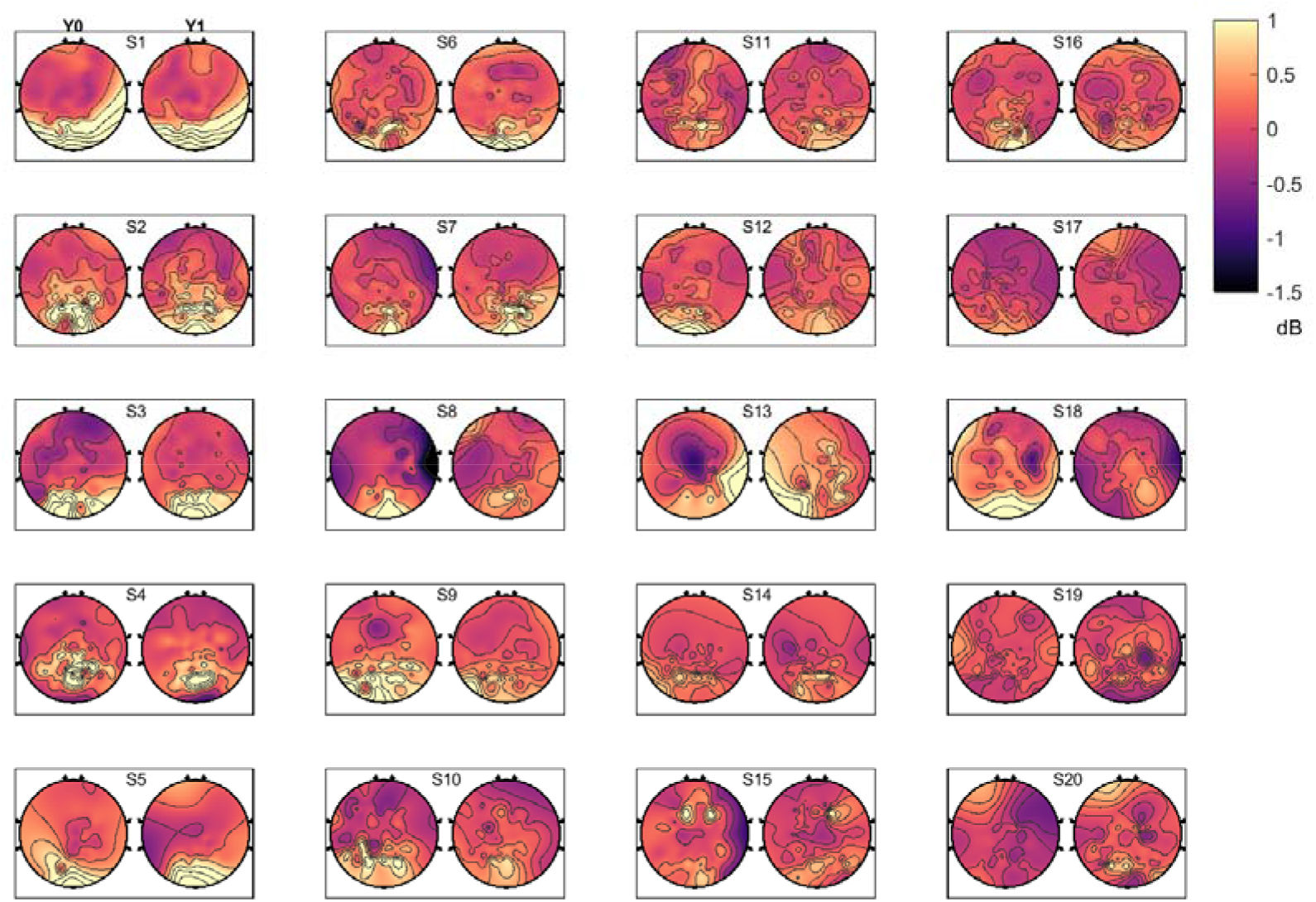
Same as Supp. Fig. 1 for 20 male subjects.

**Supp. Fig. 3.**
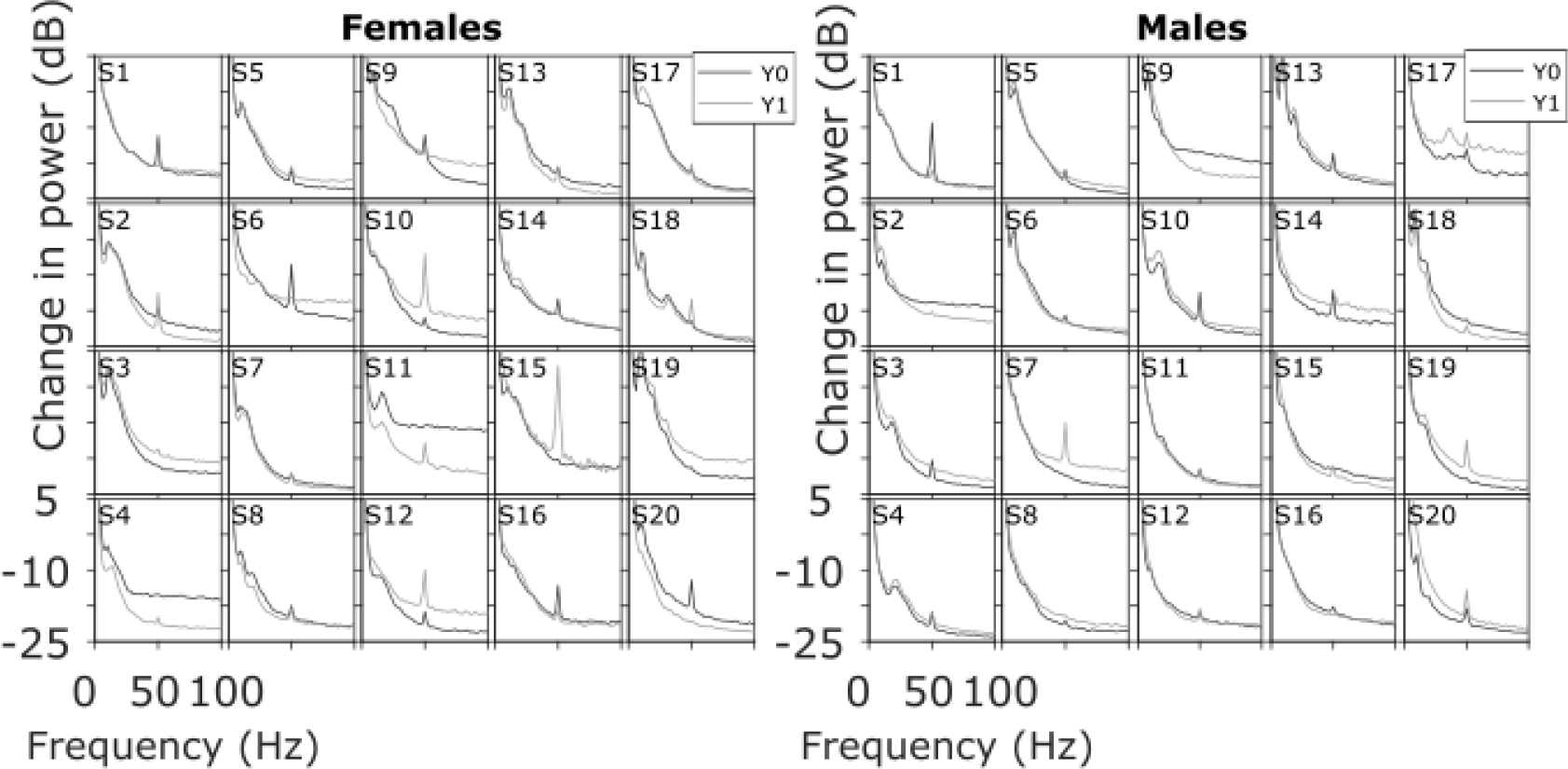
Resting state Power Spectral Density (PSD) profiles are also consistent across baseline and follow-up in males and females. Same as Fig. 3 for the absolute PSDs on the resting state data during the baseline period of the experimental data.

**Supp. Fig. 4.**
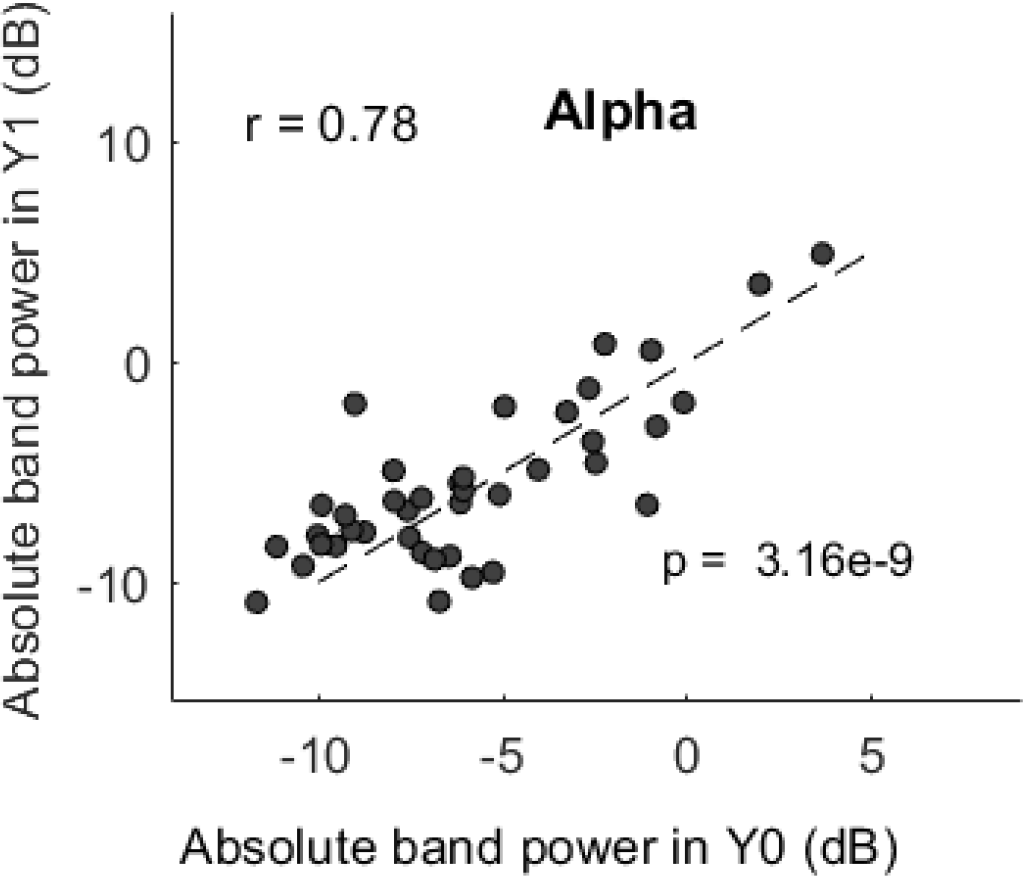
Resting state absolute alpha power is correlated across baseline and follow-up. Same as Fig. 5 for resting state absolute alpha power (computed over the baseline period).

